# Isolation of bioconcrete-producing bacteria for urea-free marine applications

**DOI:** 10.64898/2026.06.26.734652

**Authors:** Jennifer Bracewell, Farzana Nishat, Warda Ashraf, Kelli Palmer

## Abstract

Concrete repair and replacement have significant environmental and economic costs. Bacteria can form bioconcrete via microbially-induced carbonate precipitation (MICP). Bioconcrete-forming bacteria can be incorporated into concrete at mixing and heal cracks where and when they occur. Bioconcrete formation is a byproduct of alterations to the local environment occurring during normal metabolic activities of bacteria. Bacteria thus “make” bioconcrete by different metabolic mechanisms, and the environment plays a substantial role in the yield and physical properties of bioconcrete produced by a given bacterium. The ureolytic bacterium *Sporosarcina pasteurii* is a commonly used organism for MICP, but it requires urea supplementation and generates nitrogenous waste. The marine environment is understudied for bioconcrete applications, yet self-healing structures are needed in this environment, wherein urea and nitrogenous waste would be detrimental to native biota. Here, we assessed *S. pasteurii* bioconcrete production under marine-like media conditions with urea and calcium supplementation. *S. pasteurii* generated higher bioconcrete yields in these media compared to standard medium. We then designed an enrichment protocol to isolate and characterize non-urea-requiring bioconcrete-forming bacteria from Atlantic seawater. We identified isolates from the *Sulflitobacter*, *Marinobacter*, and *Bacillus* genera, two of which yielded higher bioconcrete in seawater-mimicking media compared to model non-ureolytic bacteria. Scanning electron microscopy/energy dispersive spectroscopy and Fourier transform infrared spectroscopy revealed distinct chemical and structural features of bioconcrete produced by bacteria in seawater-mimicking medium. Overall, our work establishes a pipeline for the isolation and characterization of novel bioconcrete-forming bacteria from marine samples, with application to marine self-healing materials.

**Importance:** Bacteria can stimulate the formation of calcium carbonate, referred to as bioconcrete, around their cells. Bioconcrete can be applied towards the sustainable healing of cracks in concrete structures. However, the specific chemical composition of the concrete and the environment the concrete is placed in limit the usefulness and applicability of individual bacterial species towards bioconcrete formation. Here, we focused on the marine environment, which has been less studied in the bioconcrete field relative to the terrestrial environment. We used marine-mimicking growth media to evaluate bioconcrete formation by model bacteria previously used by bioconcrete researchers as well as to isolate novel bioconcrete-forming bacteria from Atlantic seawater. Further, we used analytical chemistry and microscopy techniques to characterize the similarities and differences among the bioconcrete produced by the bacteria. Overall, we provide a framework for the isolation and characterization of bioconcrete-forming bacteria for application to sustainable infrastructure in the marine environment.

## Introduction

With the need to reduce global CO_2_ emissions, bioconcrete is increasingly researched as a sustainable alternative to manual concrete repair and replacement arising due to structure degradation via cracking. Bioconcrete, also commonly called microbially induced carbonate precipitation (MICP), is a byproduct of normal bacterial metabolic activities that produce suitable environmental conditions to precipitate calcium carbonate (1). Several bacterial metabolic activities contribute to creating optimum conditions for bioconcrete formation. Among the most studied are urease activity, carbonic anhydrase activity, and denitrification (1–3). A specific bacterium may use one or more of these as part of its normal metabolic activities. These mechanisms result in changes in the local environment around the bacterial cell that increase the local pH, calcium, and carbonate concentration, thus allowing the bacterial cell surface to act as a nucleation site for calcium carbonate, or bioconcrete, precipitation (1,4).

Urease activity is the most extensively studied mechanism for bioconcrete production, owing to its rapid and plentiful production of bioconcrete. The ureolytic mechanism of MICP starts with urea degradation by the enzyme urease, producing ammonia and carbon dioxide. Ammonia raises the local pH, promoting the formation of bicarbonate from carbon dioxide. The calcium from a calcium precursor, such as calcium acetate or calcium chloride, is attracted to the negatively charged cell surface. Together, these increase the local concentrations of calcium and bicarbonate around the cell’s surface, allowing the cell surface to act as a nucleation site for the precipitation of calcium carbonate (5).

Similar to urease activity, the carbonic anhydrase mechanism for bioconcrete production relies on an alkaline pH to precipitate calcium carbonate. Carbonic anhydrase is a widespread enzyme, with all three major classes found in prokaryotes. Carbonic anhydrases reversibly catalyze the conversion of CO_2_ to HCO_3-_, both of which can be used by the cell for metabolic or transport purposes (6). As a byproduct of these activities, bicarbonate can accumulate or be transported outside of the cell where the presence of calcium allows the precipitation of calcium carbonate.

Apart from these mechanisms, several other factors impact bioconcrete production, such as the choice of microbe and that microbe’s specific metabolic abilities, availability of nutrients/minerals, culture or growth conditions (such as temperature and oxygenation), quantity of bacteria used, and bacterial metabolic state (vegetative vs dormant/endospore) (3,7). The culture medium provides nutrients and energy sources, influencing bacterial growth. In addition, the supplementation of urea and/or calcium precursor impacts the amount and morphology of calcium carbonate produced by the bacteria (8). With sufficient levels of the appropriate nutrients, bacterial cells can actively (vegetatively) replicate. In contrast, bacterial endospores are dormant, metabolically inactive cells that can withstand extreme environmental pressures such as high heat, pressure, and desiccation that usually kill vegetative bacteria (9). The hardiness of bacterial endospores allows them to survive their incorporation into concrete during mixing and pouring. Also, the ability to remain dormant in the concrete for long periods of time ensures the bacteria remain viable when the concrete later cracks (10). Once the concrete cracks, there is an ingress of water and nutrients that allows the endospores within the crack to germinate back into a vegetative state. The resumption of normal metabolic activity facilitates bioconcrete formation and aids in the healing of cracks (11). The Gram-positive endospore-forming bacterium *Sporosarcina pasteurii* is the most commonly used model organism for bioconcrete production due to its high urease activity, which improves its bioconcrete yield, and its endospore-forming ability, which allows its incorporation into concrete mixtures (4).

The marine environment is not a commonly researched application environment for bioconcrete, although there is a need for sustainable repair of marine structures (12). The marine environment presents several challenges that do not apply to terrestrial bioconcrete. One is that the high salinity of the environment can greatly impact bacterial growth (13), especially for soil microorganisms like *S. pasteurii* (14). Another is that urea, and the ammonium resulting from urease activity, may be detrimental to marine life. This is particularly relevant for the construction of self-healing artificial coral reef structures or other structures to support marine recolonization or food production (for e.g., oyster beds). The use of bioconcrete by non-urease producing organisms may make underwater concrete structures more sustainable in the long term due to self-healing capabilities, reducing the need for human intervention for repair or replacement.

Since bioconcrete is a byproduct of normal metabolic activity, bacteria must actively use these pathways for its production. And, because different bacteria may respond differently to the same environment (for our study, the marine environment), it is important to test and compare different strains to identify the most suitable candidate for a specific environment or application. Here, we tested the ability of *S. pasteurii* to form bioconcrete under culture conditions mimicking non-marine and marine settings, all of which required the addition of urea. We then designed and implemented an enrichment culture method to isolate novel, non-urea-dependent bioconcrete-forming bacteria from seawater collected from the Atlantic coast of the United States. We identified these bacteria by whole genome sequencing and analysis and quantified their ability to form bioconcrete in a non-urea-dependent manner under marine-mimicking culture conditions. Overall, our study establishes a pipeline for the isolation and characterization of novel bioconcrete-forming bacteria from seawater.

## Materials and Methods

### Bacterial strains and culture methods

Bacteria used in this study are shown in **Table 1** along with the media used for their routine culture. Growth media used include Lysogeny Broth (LB; 10 g/L tryptone, 5 g/L yeast extract, and 10 g/L sodium chloride), ATCC medium 1376 (*Bacillus pasteurii* NH_4_-YE medium), and ATCC medium 2029 (nutrient agar, pH 9.0). For agar cultures, liquid media were supplemented with 15 g/L agar, with the exception of 20 g/L agar for ATCC medium 1376. Artificial sea medium (ASM) and artificial reef medium (ARM) were comprised of 35.96 g/L Instant Ocean Sea or Reef Salt, respectively, and 5 g/L tryptone and 1 g/L yeast extract. Sporulation medium was comprised of ASM supplemented with 5 mg/L MnSO_4_, based on a medium described previously for *B. toyonensis* spore cultivation (15). For *S. pasteurii* cultures, media were supplemented with 20 g/L urea, resulting in modified LB (LBU) and modified ASM and ARM (ASMU and ARMU). For bioconcrete quantification cultures, media were supplemented with 10 g/L calcium acetate, resulting in modified ASM and ARM (referred to here as ASMCa and ARMCa). For *S. pasteurii* bioconcerete quantification cultures, LBU, ASMU, and ARMU were supplemented with 10 g/L calcium acetate (referred to here as LBCaU, ASMCaU, and ARMCaU). The sources of the chemicals used in this study are shown in **Table S1**. A comparison of the medium components present in the base versions of the bioconcrete quantification media is shown in **Table S2**.

**Table 1.** Bacterial strains used in this study.

| Organism | Strain | Formerly classified as | Routine broth medium<sup>a</sup> | Routine agar medium<sup>a</sup> | Bioconcrete quantification media<sup>a</sup> | References for strains |
| --- | --- | --- | --- | --- | --- | --- |
| *Sporosarcina pasteurii* | ATCC 11859 | *Bacillus pasteurii* | LBU | ATCC medium 1376 | LBCaU, ASMCaU, ARMCaU | 51 |
| *Bacillus toyonensis* | UTDF19-29B | - | LB | ASM | ASMCa | 52 |
| *Alkalihalophilus pseudofirmus* | ATCC 700159 | *Bacillus pseudofirmus*; *Bacillus firmus* | ASM | ATCC medium 2029 | ASMCa | 30, 53 |
| *Marinobacter* sp. | SBS5 | - | ASM | ASM | ASMCa | This work |
| *Sulfitobacter* sp. | SBS6 | - | ASM | ASM | ASMCa | This work |
| *Bacillus* sp. | SBS7 | - | ASM | ASM | ASMCa | This work |

<sup>a</sup>See the Materials and Methods section for media compositions and naming conventions.

For modified LB media (LBCaU or LBU), the base LB medium was made as a separate 2X solution and sterilized by autoclaving at 121°C for 20 minutes. Solutions containing urea and/or calcium acetate in water were made at 2X concentrations and filter sterilized by vacuum filtration with a 0.22 μm filter to avoid precipitation after autoclaving. The two 2X solutions were then aseptically combined, resulting in sterile liquid media for use. The modified artificial sea and reef media and the sporulation medium were filter sterilized to achieve sterility. All other media were sterilized by autoclaving at 121°C for 20 minutes.

Agar cultures were incubated at room temperature at ∼21°C. Routine overnight broth cultures were inoculated with a single colony each from a fresh streak plate. Broth cultures were incubated at 30°C with shaking at 150 rpm. Bacterial cultures were cryopreserved by mixing 750 μL sterile 50% glycerol with 750 μL of overnight broth culture with storage at -80°C.

### Bacterial isolation from seawater

Seawater was collected in sterile 50 mL conical tubes on June 9^th^, 2023 at GPS coordinates 41.370941, -71.497279 and shipped on ice to Richardson, TX. Once received, cryopreserved freezer stocks were created as described above, using seawater as the “culture”. To screen for bacteria that produce bioconcrete, the stocks then were used to inoculate ARM broth cultures which were then used as inocula for 15 mL ARMCa bioconcrete quantification cultures. After 15 days of incubation, cultures with visually observable bioconcrete production were cryopreserved as above, and the remaining culture was used for bioconcrete quantification. The population that produced the most bioconcrete (by dried precipitation weight) was streaked on ASM agar for single colony isolation, yielding 6 isolates, which were cryopreserved as above.

### Gram staining

Gram staining was performed using the BD Gram Stain Kit and following the protocol published by the American Society for Microbiology (16).

### Sporulation test

Isolates were cultured overnight in ASM, pelleted, resuspended in sporulation medium, and cultured for one week to test sporulation capabilities. Ability to sporulate was initially evaluated by phase-contrast microscopy by examining cultures for phase-bright spots. Endospores were visualized under bright field microscopy after staining using the Shaeffer-Fulton method (17), with modifications as previously recommended (18). Briefly, a few microliters of liquid culture were dried on a glass slide before heat fixing. The slides were put in an empty Petri dish, with a small square of paper towel placed over the slide. The paper towel was saturated with malachite green solution (0.25 mg/mL), and the lid was put on the Petri dish. The closed dish was left to saturate at room temperature for 30 minutes. Then, the slide was rinsed with water, counterstained 30 seconds with safranin, and rinsed again. The slide was visualized after drying.

### DNA isolation and genome sequencing

DNA was isolated from overnight bacterial cultures as performed previously (19), except that 80 mg/mL lysozyme was used. Illumina sequencing was performed by Seqcenter (Pittsburgh, PA). Illumina reads were assembled into contigs using the De Novo Assembly tool on CLC Genomics Workbench with default parameters. Genome assemblies were annotated using RAST (20–22), and an annotated 16S rRNA gene was used for preliminary taxonomic assignment using NCBI BLAST to identify the closest species. Among the isolates, 3 species were identified, and one isolate of each species was submitted to NCBI for genome sequence archiving, additional taxonomic analysis, and annotation. The Whole Genome Shotgun project for *Marinobacter* sp. SBS5 has been deposited at DDBJ/ENA/GenBank under the accession JBLWAZ000000000. The Whole Genome Shotgun project for *Sulfitobacter* sp. SBS6 has been deposited at DDBJ/ENA/GenBank under the accession JBLWBA000000000. The Whole Genome Shotgun project for *Bacillus* sp. SBS7 has been deposited at DDBJ/ENA/GenBank under the accession JBLWBB000000000.

### Bioconcrete quantification protocol

Bioconcrete production by bacterial cultures was quantified at various timepoints over 16 days (*S. pasteurii* at days 1, 6, and 16; *B. toyonensis*, *Ahb. pseudofirmus* and the sea isolates at day 16) using a protocol modified from Reeksting et al. (23). All quantifications were performed in three independent biological replicates. A step-by-step numbered version of this protocol is shown in **Supplemental Text S1**.

Overnight cultures were grown in the routine medium specified in **Table 1** for the specific bacterium. The optical density of the culture was measured at 600 nm (OD_600_) using a Thermo Scientific GENESYS 30 Visible Spectrophotometer. These readings were used to calculate the volume of the overnight culture required to inoculate the bioconcrete quantification medium to a starting OD_600_ of 0.001. For *S. pasteurii*, 50 mL of bioconcrete quantification medium was inoculated with the calculated amount of the overnight culture and incubated. After 2 hours, 15 mL culture was aliquoted into 50 mL centrifuge tubes. Since the other bacterial species in this study were analyzed at only one timepoint, this step was omitted for them. Instead, 15 mL of quantification medium was inoculated directly in a 50 mL centrifuge tube. For all bacteria, the lids of the 50 mL tubes were taped in place one half turn from being completely shut to allow consistent oxygenation throughout the experiment. The tubes were incubated until their timepoint for quantification. A tube was removed for quantification at the timepoint 1, 6, and/or 16 days after inoculation and vortexed thoroughly. The pH was measured by dripping 30 μL of culture onto pH strips (Fisher Brand). Serial dilutions were performed (1:10 dilutions in sterile phosphate buffered saline) to calculate the viable cell count, or colony forming units (CFU)/mL, remaining in the culture. These dilutions were plated using the track dilution method described by Jett et al. (24). The colonies were typically counted after 3 days to calculate CFU/mL in the bioconcrete culture. The rest of the culture, still in the 50 mL centrifuge tube, was then centrifuged at 980 x g for 4 minutes (Sorvall RC 6+ Centrifuge with F13S-14x50cy rotor), and the supernatant was aspirated. The pellet was washed with 40 mL of autoclaved deionized water, resuspended by vortexing the pellet, recentrifuged, and the supernatant was aspirated. The pellet was washed in this manner a total of three times. After washing, the pellet was resuspended in 1 mL autoclaved deionized water and transferred using a wide-bore pipette tip to a pre-weighed 1.5 mL microcentrifuge tube to dry. The tube was left open but covered with a Kim Wipe (Science Brand) to allow for evaporation and dried at 55°C for ≥2 days. Finally, the Kim Wipe was removed, and the tube with the dry bioconcrete was weighed. The weight of the empty tube was subtracted to yield the weight of the precipitated bioconcrete. Each experiment was paired with a control tube of uninoculated media that underwent the same protocol to monitor for contamination or measurable abiotic precipitation during the experiment.

### SEM/EDS analysis of collected bioconcrete

Scanning electron microscopy (SEM) was done on a Zeiss EVO LS 15 at the University of Texas at Dallas Imaging Core. Energy dispersive spectroscopy (EDS) was performed using an Oxford EDS detector at the University of Texas at Dallas Imaging Core. Bioconcrete samples were affixed to 12 mm aluminum stubs (Electron Microscopy Sciences, 12.7 mm diameter, 3 mm pin, 12 mm total high, cat #75500) using double-sided carbon tape (Electron Microscopy Sciences, 8 mm wide, cat #77816). To prepare each sample, a small dash of the dried bioconcrete powder was placed on a piece of parafilm and then resuspended in water using a micropipette which was also used to transfer the resuspended sample to double-sided carbon tape that was already affixed to the aluminum stub. Prior to imaging, the prepared samples were kept at 75°C for ≥2 hours to dry and to reduce outgassing while under vacuum.

### Fourier Transform Infrared (FTIR) spectroscopy

FTIR was performed using a Nicolet iS5 spectrometer operating in Attenuated Total Reflection (ATR) mode. For each specimen, 32 scans were collected at a spectral resolution of 4 cm⁻¹ over the wavenumber range of 400–4000 cm⁻¹. The signal-to-noise ratio was maintained below 3:1 throughout the measurements.

## Results

### Establishing the protocol used for bioconcrete quantification in this study

We based our bioconcrete quantification protocol around one previously published by Reeksting et al. (23). We chose *S. pasteurii* as the reference organism for bioconcrete quantification because it is one of the best studied bioconcrete-producing organisms due to its alkalophilicity, high urease activity, and ability to form bacterial spores (4). We began our study by converting the protocol published by Reeksting et al. (23), to numbered form and performing bioconcrete precipitation assays per that protocol. Our results are shown in **Figure S1** and **Dataset S1**; Dataset S1 shows raw bioconcrete precipitation weights, pH values, and CFU/mL values for all independent experimental replicates. We did not achieve the ∼6 g/L bioconcrete precipitation weight expected for *S. pasteruii* culture in LBCaU medium, as obtained by Reeksting et al. (23), instead achieving only ∼3-4 g/L, and our bioconcrete yields were also variable across independent experimental replicates. We then made changes to the protocol to improve yield and reproducibility in our hands (**Supplemental Text S1** and see Discussion for consideration of factors that contributed to yield and variability). Using this modified protocol, we achieved the expected amount of bioconcrete precipitation for *S. pasteurii*, with low variability across independent experimental replicates (**Figure S1)**. The modified protocol was implemented for all following bioconcrete experiments in this study.

### Robust production of compositionally distinct bioconcrete by *S. pasteurii* in media mimicking seawater and reef, supplemented with urea and calcium

In the artificial sea and reef media supplemented with urea and calcium (ASMCaU and ARMCaU, respectively), bioconcrete yields from *S. pasteurii* at the day 16 timepoint were at least 1.5 g/L higher than those obtained with modified LB medium (**Figure 1A)**. Because *S. pasteurii* produced more bioconcrete in ASMCaU than in modified lab medium, it could be a promising bacterium for special-use marine or other high salt applications where the inclusion of urea in the concrete mix and the release of urease waste products is environmentally acceptable.

**Figure 1:**
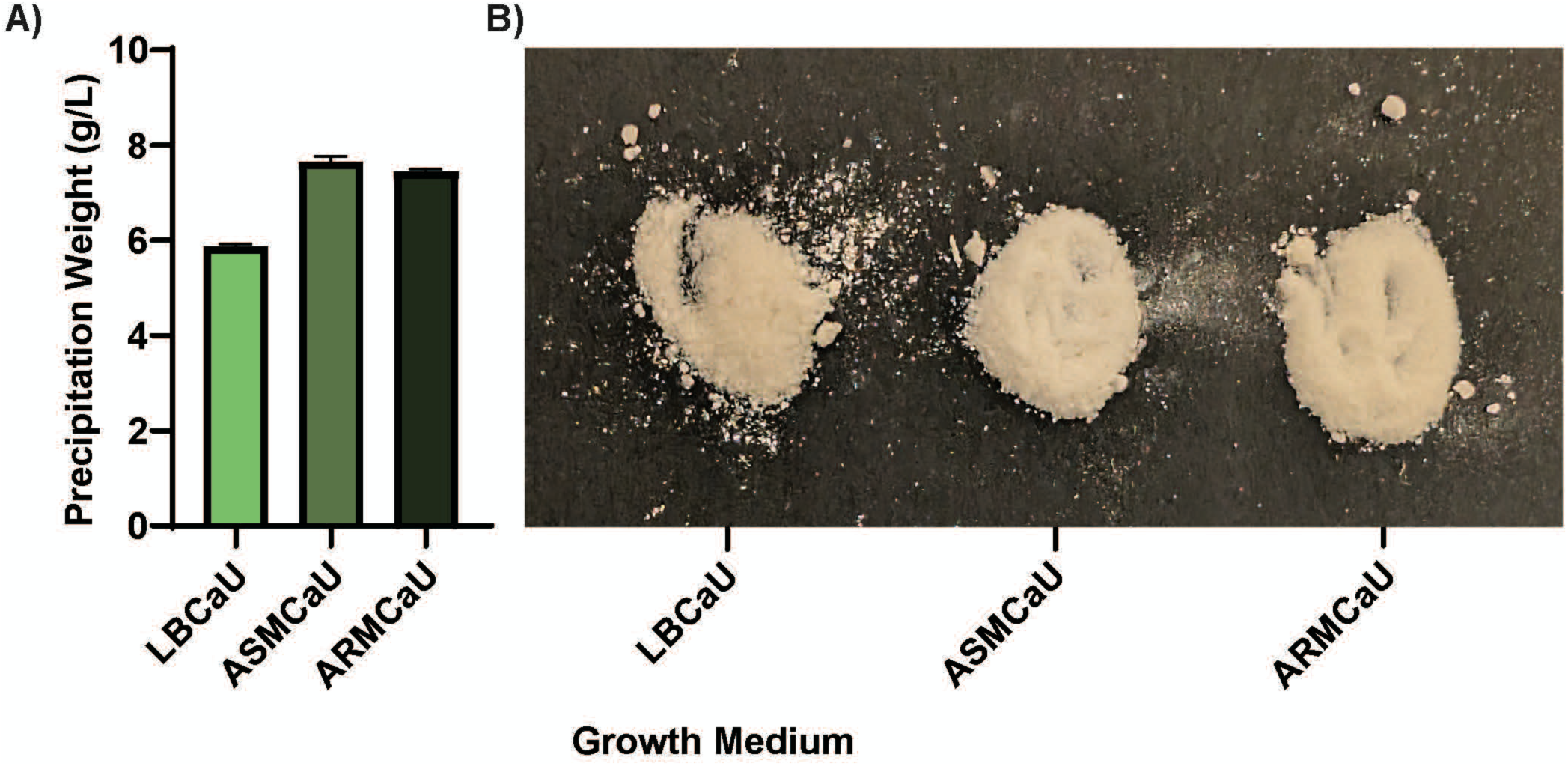
Bioconcrete precipitated by S. *pasteurii* in various growth media. A) Amount of bioconcrete precipitated after 16 days by S. *pasteurii* in LBCaU, ASMCaU, and ARMCaU using the revised protocol. B) Image of the dried bioconcrete produced by S. *pasteurii* in LBCaU, ASMCaU, and ARMCaU respectively.

**Figure 1B** shows bioconcrete collected from the *S. pasteurii* cultures. We observed changes in the color and coarseness of the bioconcrete grains in the macro scale when different culture media were used. SEM analysis shows that the bioconcrete was mainly spherical and agglomerated in morphology, with a range from a few micrometers to ≥20 micrometers (**Figure 2**). The EDS results of the samples verified the bioconcrete to be calcium carbonate (**Figure 2**). A difference in morphology was observed in the bioconcrete precipitated in ASMCaU and ARMCaU compared to each other and to the standard lab medium (**Figure 2**, panels C, E, and A, respectively). Compared to the very smooth, spherical, bioconcrete formed in the modified lab medium, the bioconcrete formed in the ASMCaU showed much smaller and less rounded spherical grains that were still very clustered (**Figure 2C**), while the grains precipitated in the ARMCaU showed more flaky, agglomerated spheres (**Figure 2E**). Of note, the bioconcrete precipitated in ASMCaU and ARMCaU had small amounts (2.9-4.4%) of magnesium detected by EDS. The presence of magnesium may impact structurally relevant properties of the bioconcrete, which should be evaluated in the future.

**Figure 2:**
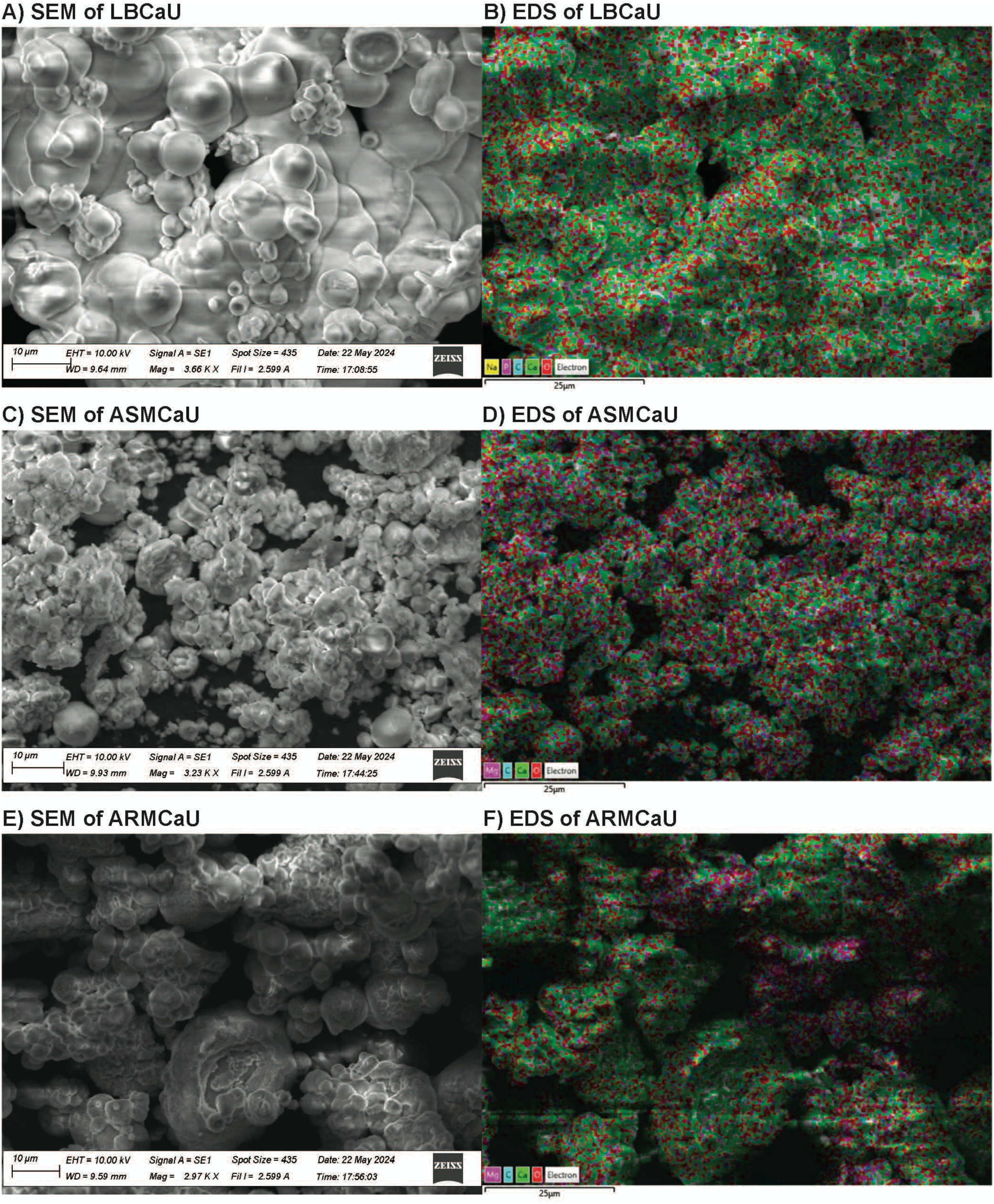
SEM images of the bioconcrete produced in the S. *pasteurii* quantification experiments after 16 days, and their corresponding EDS false color images. In the EDS images, blue indicates carbon, green indicates calcium, and red indicates oxygen. A) LBCaU. B) EDS of A. C) ASMCaU. D) EDS of C. E) ARMCaU. F) EDS of E.

**Figure 3** presents the FTIR spectra of bioconcrete produced by *S. pasteurii* cultivated in the different media. Carbonate ions exhibit four characteristic vibrational modes: symmetric stretching (*ν_1_*) near 1080 cm^-1^, out-of-plane bending (*ν_2_*) between 850 and 880 cm^-1^, asymmetric stretching (*ν_3_*) within the range of 1400–1500 cm^-1^, and in-plane bending (*ν_4_*) between 690 and 760 cm^-1^ (25,26). Among these vibrational modes, the *ν_4_* region is particularly useful for distinguishing calcium carbonate polymorphs as calcite, vaterite, and aragonite exhibit characteristic absorption bands in this region, whereas significant peak overlap occurs in the other vibrational modes.

**Figure 3:**
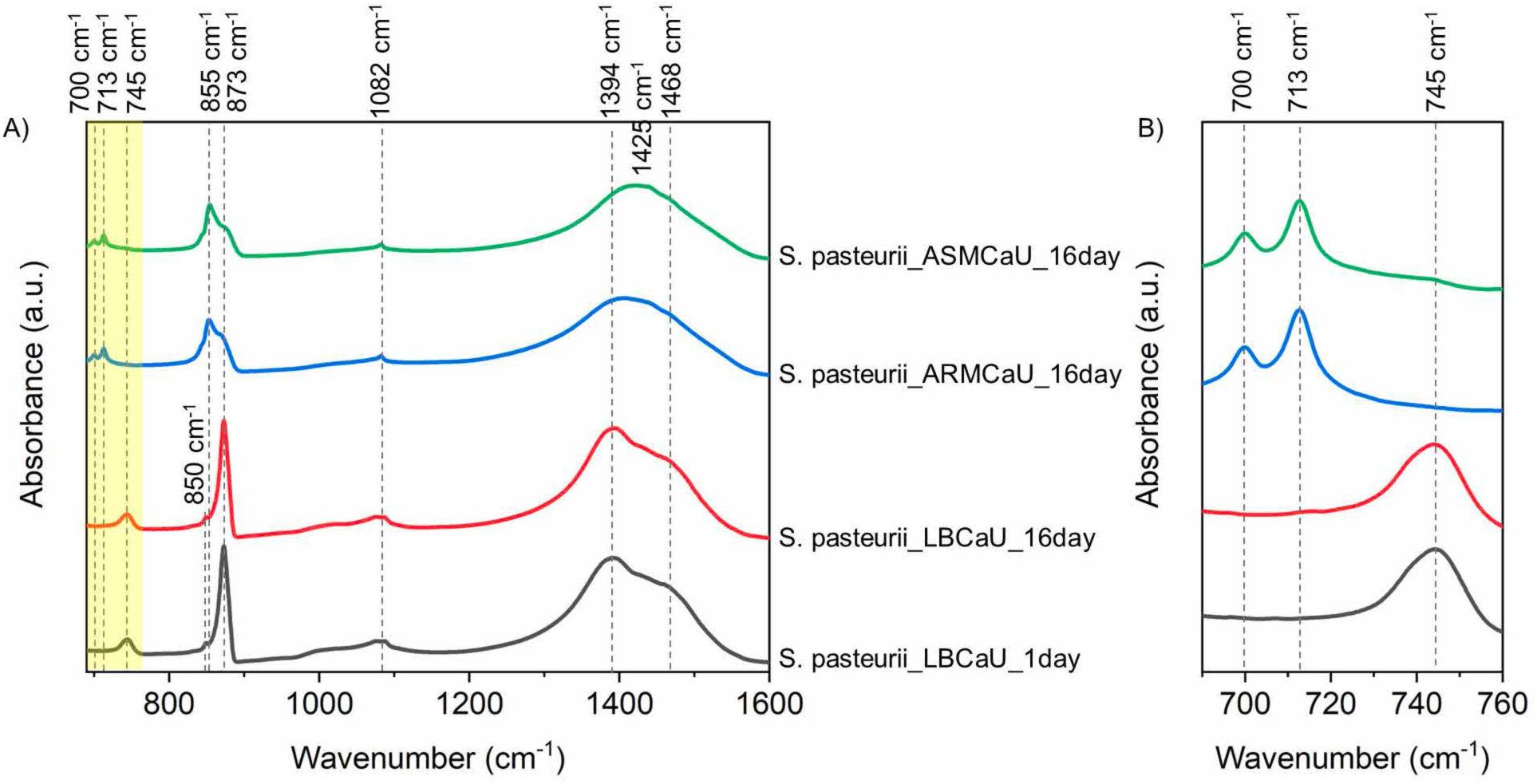
FTIR spectra of bioconcrete produced by ureolytic bacteria (S. *pastuerii)* in different media: A) shown from range 690-1600 cm-^1^, **B)** shown from range 690-760 cm-^1^.

A broad absorption band observed between 1400 and 1500 cm^-1^ in all *S. pasteurii*-produced bioconcrete samples suggests the presence of metastable calcium carbonate phases, including vaterite and amorphous calcium carbonate. To further identify the carbonate polymorphs, the *ν_4_* region (690–760 cm^-1^) was examined in detail (**Figure 3B**). Calcite is typically characterized by an absorption band near 713 cm^-1^, whereas vaterite exhibits a characteristic peak around 745 cm^-1^. Aragonite can be identified by the simultaneous presence of bands near 700 and 713 cm^-1^ (25,26).

The bioconcrete produced by *S. pasteurii* in LBCaU medium exhibited a broad absorption band approximately at 745 cm^-1^, indicating the predominance of vaterite. This spectral feature was detected from day 1 and remained largely unchanged throughout the 16-day incubation period. In contrast, samples produced in ASMCaU and ARMCaU media displayed two distinct absorption bands at approximately 700 and 713 cm^-1^, confirming the formation of aragonite. The presence of calcite cannot be ruled out in these samples because its characteristic peak overlaps with the aragonite band at 713 cm^-1^.

Overall, these results verify that marine-relevant growth media impact the yield, morphology and composition of bioconcrete produced by *S. pasteurii*.

### Enrichment culture method to identify bacterial candidates for urea-free marine bioconcrete applications

Urea supplementation and urease activity for bioconcrete production increases nitrogen levels, which is detrimental to marine environments (27,28). Additionally, using bacteria native to the environment where bioconcrete will be implemented may help ensure that the bacteria are adapted to and will be metabolically active in that environment. To identify organisms suitable for urea-free marine bioconcrete applications, we designed an enrichment culture strategy using seawater as inoculum. As depicted in **Figure 4**, to enrich for bioconcrete-forming organisms, we first quantified the bioconcrete produced by cultures inoculated with seawater. These bioconcrete-producing consortia were cryopreserved after a 15 day enrichment, and the preserved stock that had produced the most bioconcrete by weight during the experiment was then struck to isolation. By selecting the highest performing consortia to isolate bacteria from, we narrowed down the number of isolates to where we could easily test their independent bioconcrete production in pure cultures.

**Figure 4:**
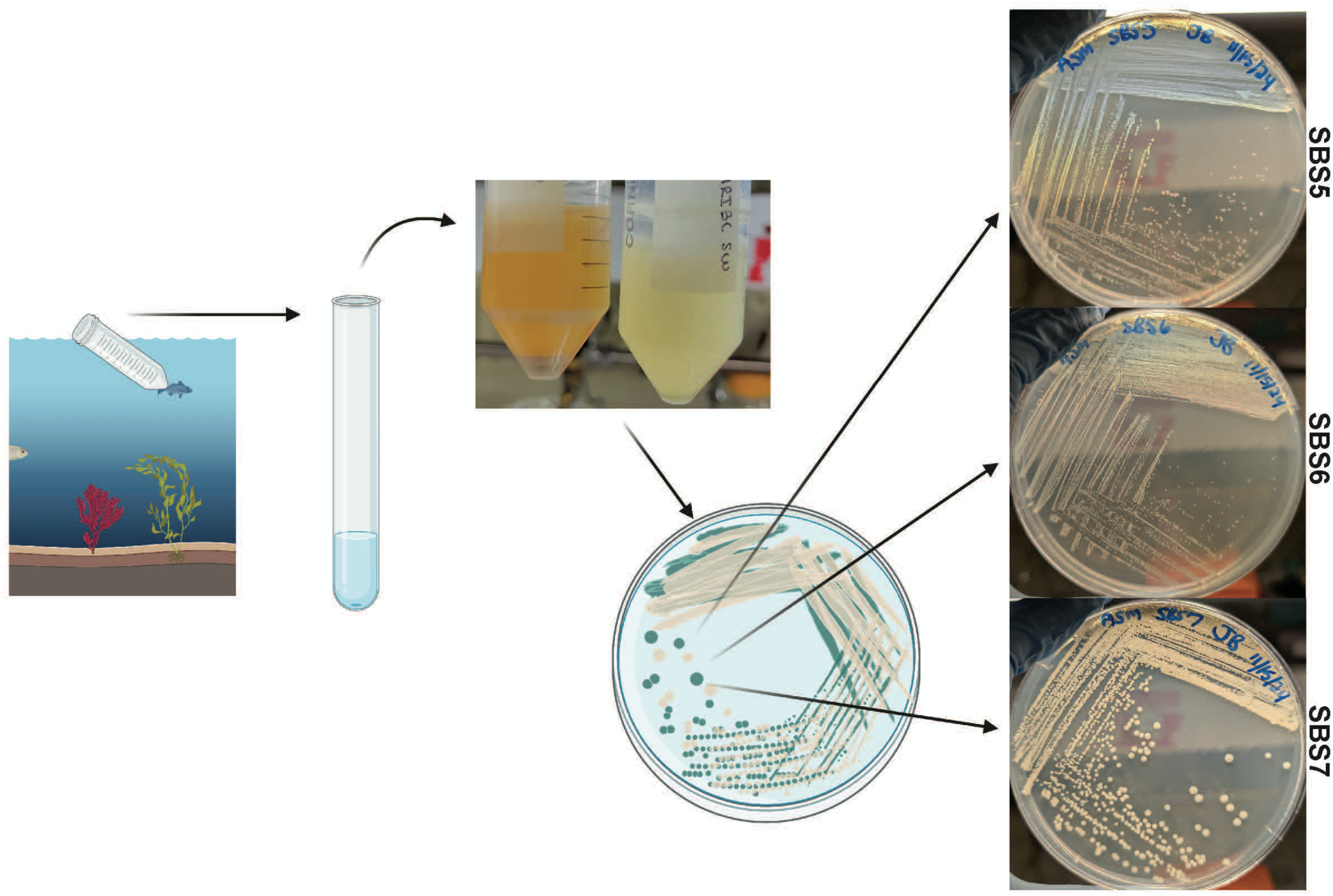
Isolation of bioconcrete forming bacteria from sea water. Sea water was collected, screened for bioconcrete production in ARMCa (example cultures pictured top middle), and then streaked to isolated species (the three named sea isolates’ streaks pictured on the right). Figure created in BioRender.com.

We used genomic DNA sequencing to obtain preliminary species assignments for the 6 isolates obtained from the highest performing enrichment culture. Analysis of the 16S rRNA genes of the sea isolates identified by initial genome annotations (by RAST; 20–22) revealed there to be 3 different species among the isolates, with the closest relatives being *Sulfitobacter pontiacus* (two of the isolates, both with 99.79% identity to members of this species, one called *S. sp.* SBS6 hereafter), *Marinobacter litoralis* (three of the isolates, with 99.80, 99.87, and 99.87% identities, one called *M. sp.* SBS5 hereafter), and *Bacillus sanguinis* (one of the isolates, with 100% identity, called *B. sp.* SBS7 hereafter). Genome sequences for the three named isolates (one representing each species) were submitted to Genbank for formal annotation and species verification. **Table S3** shows the assembly quantity, contig number, and other features of the genome assemblies. The closest relatives for these isolates were identified with higher resolution by percent average nucleotide identity comparison with genome sequences of type strains as part of the Genbank automated pipeline (**Table 2**). While *Bacillus* species are commonly associated with MICP and bioconcrete production, to our knowledge, the *Marinobacter* and *Sulfitobacter* genera have not.

**Table 2.** Characteristics of the three seawater isolates used for bioconcrete quantification studies.

| **Isolate:** | ***Marinobacter sp.***<br>**SBS5** | ***Sulfitobacter sp.***<br>**SBS6** | ***Bacillus sp.* SBS7** |
| --- | --- | --- | --- |
| NCBI Best Match<sup>a</sup> | *Marinobacter litoralis*<br>(87.41%) | *Sulfitobacter pontiacus* (97.54%) | *Bacillus albus*<br>(98.01%) |
| 16S rRNA sequence identity<sup>b</sup> | *Marinobacter litoralis* | *Sulfitobacter pontiacus* | *Bacillus sanguinis* |
| Cellular morphology | Rods | Cocci, some with filamentous extrusions | Rods |
| Gram staining result | Gram-negative | Gram-negative | Gram-positive |
| Endospore formation | No | No | Yes |

<sup>a</sup>Best taxonomic match based on whole genome average nucleotide identity (ANI) analysis performed by NCBI after automated genome annotation. The percent ANI for the best match type-strain is shown in parentheses .
<sup>b</sup>Highest nucleotide identity match identified by NCBI BLASTn analysis.

The three isolates were additionally evaluated by Gram staining and cell morphological analysis using phase contrast microscopy (**Table 2**). Of our three isolates, SBS7 is the only endospore-former as verified by sporulation in sporulation medium. Endospores from SBS7 were visible under bright field microscopy after staining using the Shaeffer-Fulton method (17) and were phase-bright under phase contrast microscopy (**Figure 5**).

**Figure 5:**
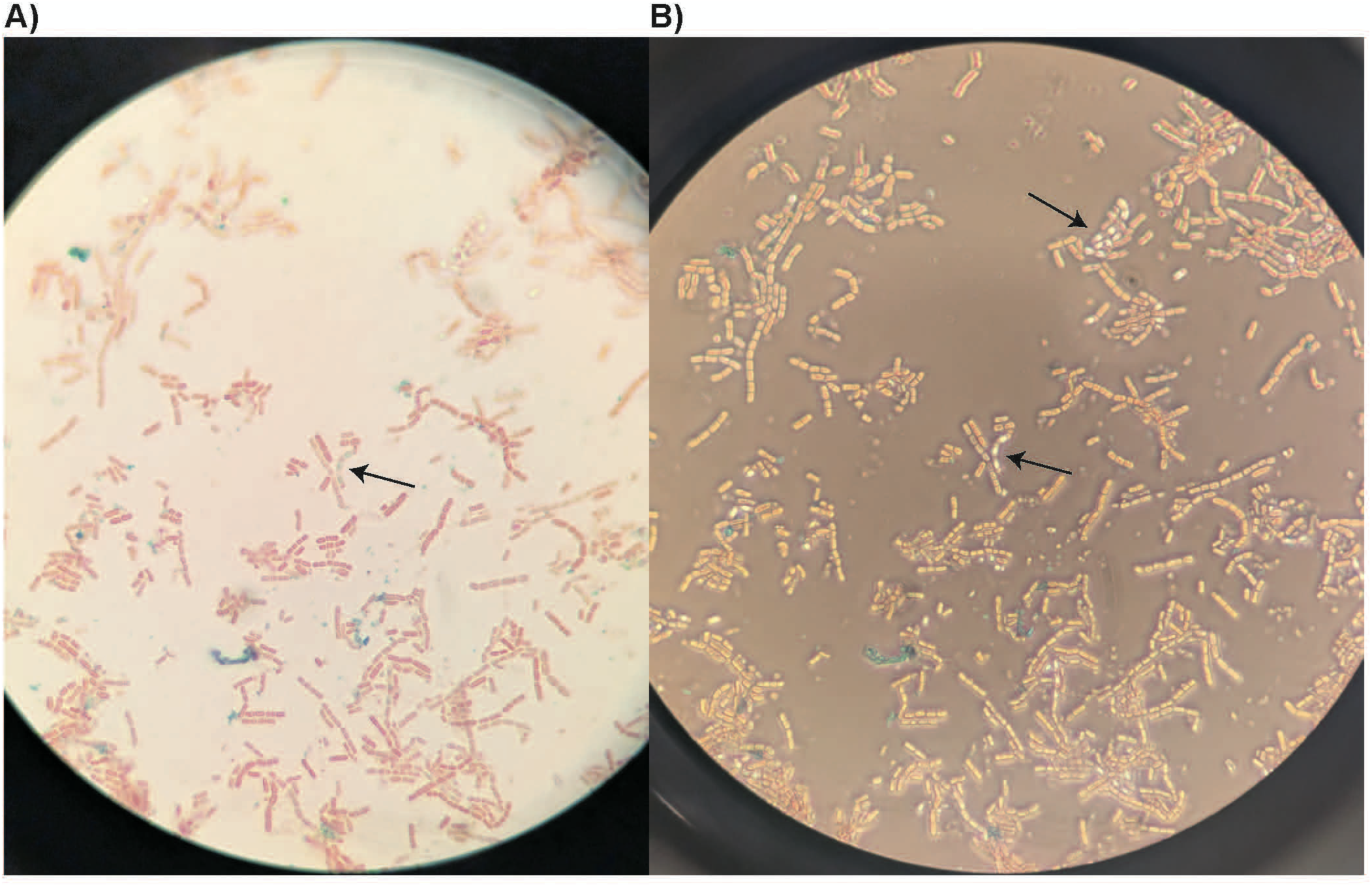
Images of *B. sp.* S8S7 at 1000X magnification under A) light microscopy after spore staining (Shaeffer-Fulton’s method) and B) the same slide viewed with phase contrast. Arrows indicate endospores, which are blue/green or phase bright in the respective images.

### Bioconcrete precipitation by non-ureolytic strains in ASM supplemented with calcium

We compared the bioconcrete precipitation yielded by the three seawater isolates and two additional non-ureolytic strains, each cultured in ASM supplemented with calcium (ASMCa) for 16 days. For the additional strains, we chose *Alkalihalophilus pseudofirmus* because it is a commonly studied nonureolytic bioconcrete-forming organism (3,29) that is also alkalophilic, halophilic, and forms endospores (30). We also chose *B. toyonensis,* which has been isolated from the sea environment (15) and is also non-ureolytic, alkalitolerant, halotolerant, and endospore-forming (31). We compared these bioconcrete yields with that produced by *S. pasteurii* in the same medium for the same time, with the exception that urea was added to the medium (note that the *S. pastuerii* data from **Figure 1** are replotted in **Figure 6** for ease of comparison). As previously reported (23,32), we observed that *S. pastuerii* could not be cultured in medium lacking urea, except in the case of media holdover, as reported by Ma et al. in 2020 (32), which allows for slight growth in media without urea.

**Figure 6:**
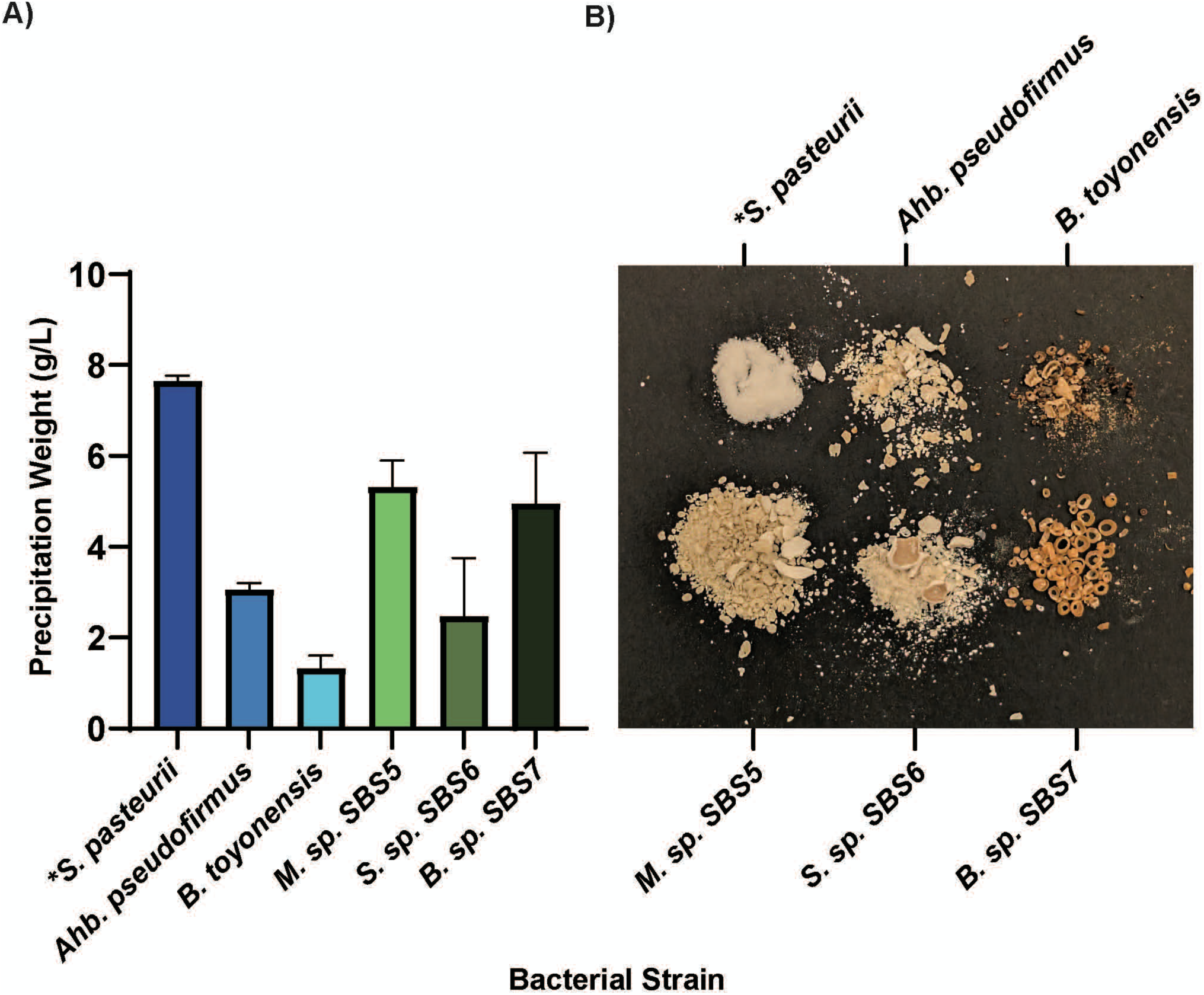
Bioconcrete precipitated by various bacteria. A) Amount of bioconcrete precipitated by S. *pasteurii, B. toyonensis, Ahb. pseudofirmus,* and the three sea isolates after 16 days in modified ASM. B) Image of the dried bioconcrete produced by the different bacterial strains in *ASMCa media. Top left to right is S. *pasteurii* (*in ASMCaU), *Ahb. pseudofirmus,* and *B. toyonensis.* Bottom left to right is *M. sp.* S8S5, S. *sp.* S8S6, and *B. sp.* S8S7. Different species produce bioconcrete in the absence of urea.

The other bacteria tested for bioconcrete production in ASMCa did not produce as much bioconcrete yield as *S. pasteurii* after 16 days, but two of the sea isolates, *M. sp.* SBS5 and *B. sp.* SBS7, performed better than *Ahb. pseudofirmus* and *B. toyonensis* in the artificial sea medium, as can be seen in **Figure 6A**. In the urea-free cultures, the pH increased to 7-8 while the control media stayed at pH 6-7. The calculated CFU/mL for each bacterium plated at day 16 of the experiments showed cell counts of ∼10^4^ for *Ahb. pseudofirmus*, ∼10^5^ for *S. pasteurii*, *B. toyonensis* and SBS6, ∼10^6^ for SBS5, and ∼10^7^ for SBS7 (**Dataset S1**), indicating enhanced survival of SBS7 in bioconcrete cultures relative to the other endospore-forming bacteria in this study.

A representative image of the bioconcrete produced by the non-ureolytic strains is presented in **Figure 6B**. The bioconcrete produced by *S. pasteurii* in the same media but with added urea is also shown for ease of comparison. This image showcases the wide range of bioconcrete morphologies that different bacterial strains can produce. SEM analysis of this bioconcrete is shown in **Figure 7** (again with the comparable *S. pasteurii* data for ease of comparison). In general, the bioconcrete produced by the non-ureolytic strains is much coarser and in larger “chunks” than that produced by *S. pasteurii*. EDS analysis verified that this bioconcrete is calcium carbonate. The EDS results also show much lower amounts of magnesium was incorporated into the non-ureolytically produced bioconcrete than the bioconcrete produced by *S. pasteurii* (**Dataset S1**).

**Figure 7:**
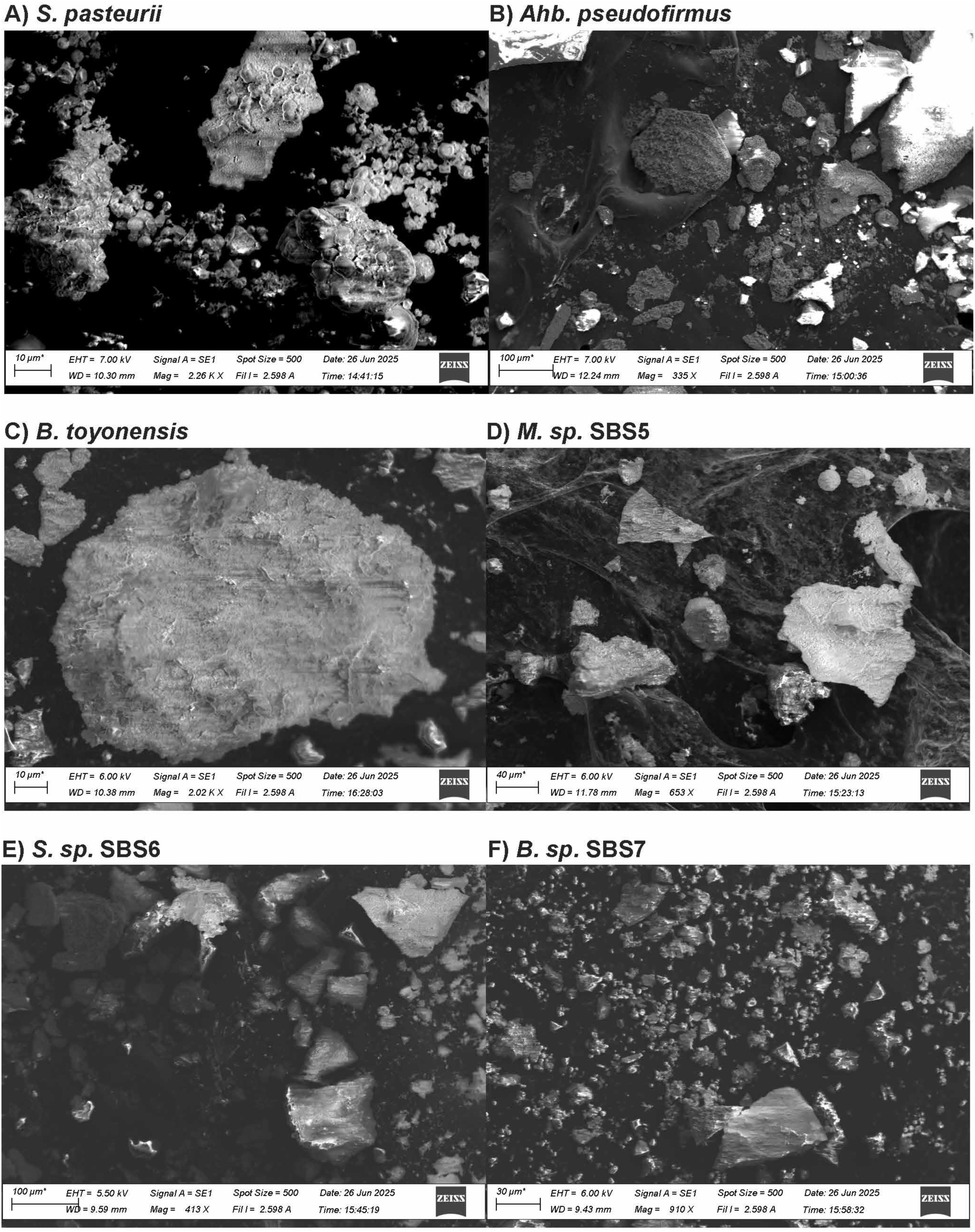
SEM images of the bioconcrete produced by the different bacterial species in ASMCa after 16 days. The bioconcrete collected from the non-ureolytic strains tends to be coarser and larger than that produced by S. *pasteurii.* A) S. *pasteurii* in ASMCaU, the same sample as Figure 2E imaged on a different day. B) *Ahb. pseudofirmus.* C) *B. toyonensis.* D) *M. sp.* S8S5. E) S. *sp.* S8S6. F) *B. sp.* S8S7.

**Figure 8** shows the FTIR spectra of bioconcrete produced by non-ureolytic bacterial strains cultured in ASMCa medium after 16 days of incubation. Similar to the samples produced by *S. pasteurii*, a broad absorption band was observed in the 1400–1500 cm^-1^ region, indicating the presence of metastable calcium carbonate phases, with clear split peaks observed for *Ahb. pseudofirmus* and *S. sp.* SBS6.

**Figure 8:**
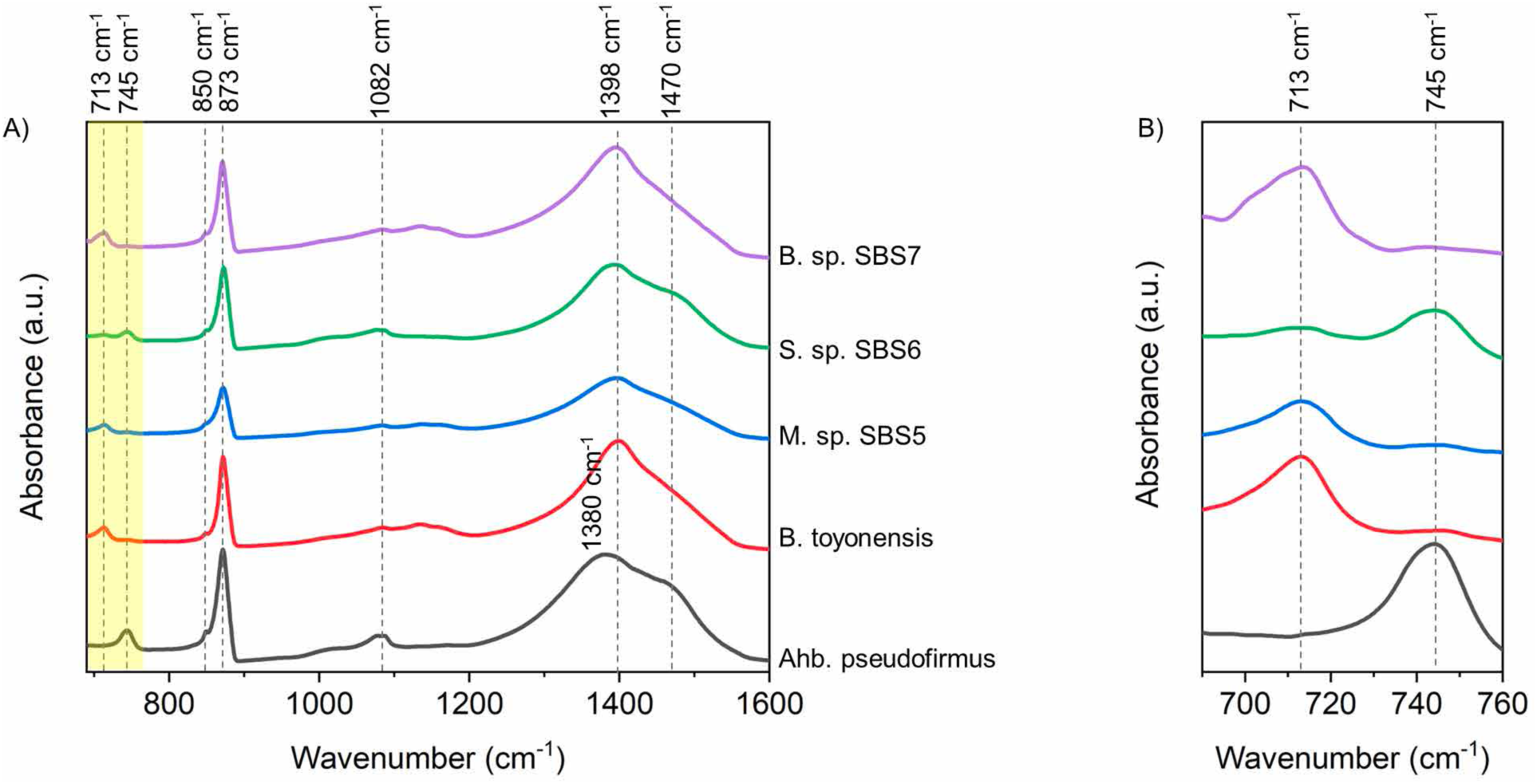
FTIR spectra of bio-concrete produced by non-ureolytic bacteria in ASMCa at day 16: A) shown from range 690-1600 cm-^1^, **B)** shown from range 690-760 cm-^1^.

Detailed investigation of the *ν_4_* region (690–760 cm^-1^) revealed notable differences among the bacterial strains (**Figure 8B**). Bioconcrete produced by *Abh. pseudofirmus* exhibited a broad peak around 745 cm⁻¹, indicating the presence of vaterite. A similar spectral feature was observed for *S. sp.* SBS6; however, a minor shoulder near 713 cm^-1^ suggested the coexistence of a small amount of calcite. In contrast, bioconcrete produced by *B. toyonensis*, *M. sp.* SBS5, and *B. sp.* SBS7 displayed a dominant absorption band near 713 cm⁻¹, characteristic peak of calcite, accompanied by a weak shoulder around 745 cm^-1^ attributable to vaterite.

A notable distinction between ureolytic and non-ureolytic bacterial systems was the absence of aragonite-specific absorption bands in the bioconcrete produced by non-ureolytic bacteria. Unlike the *S. pasteurii* samples, no clear evidence of aragonite formation was detected in these specimens.

Altogether, these results show that morphology and composition of bioconcrete varies with the bacterial species and their growth conditions.

## Discussion

In this study, we analyzed bioconcrete production by model and “wild” bioconcrete-forming bacteria in culture media mimicking the sea and reef environments. Our goal was to identify bacteria that could effectively produce bioconcrete in a urea-independent manner, thus we designed and employed an enrichment culture-based protocol to identify promising candidates from seawater.

Our initial step in this work was to reproduce a previously reported method for bioconcrete quantification (23). That prior work also utilized *S. pasteurii* as a control, establishing a baseline by which we could compare bioconcrete yields across their group and ours. We believe there were several steps that resulted in a loss of bioconcrete or that introduced variation among replicates in our hands; to address these, we added modifications to the protocol. First, the amount of bacteria used to inoculate the quantification cultures significantly influences the bioconcrete yield (33). We thus standardized our inoculation of bioconcrete cultures to an OD_600nm_ to 0.001, which likely helped lower the variability we initially observed across biological replicates. Second, the use of glass flasks for culturing introduced a point of loss, as a substantial quantity of bioconcrete produced during the protocol adhered to the walls and bottom of the flask and thus could not be fully quantified. Even thoroughly mixing the culture at quantification timepoints did not remove the adhered bioconcrete; it required scraping off for complete recovery. Thus, we utilized polypropylene 50 mL conicals for bioconcrete cultures, with an individual conical for each timepoint to be analyzed (i.e., no repeat sampling from a single culture vessel). Also, we observed that centrifugation for 2 minutes was insufficient to fully form the bioconcrete pellet, especially for cultures that make finer grains. Removing supernatant prior to fully forming the pellet also caused a loss in the amount of bioconcrete able to be quantified, and this loss differed by the morphology of the bioconcrete itself. Increasing the centrifugation time to 4 minutes helped achieve a more stable pellet for washing. Notably, centrifuged bioconcrete does not form a settled-enough pellet for typical decanting. We removed the supernatant instead by vacuum aspiration to prevent losing some of the bioconcrete pellet during each wash step. Finally, due to the high static charge, the dried bioconcrete was resuspended in water then transferred to the vessel that it was weighed in before drying. This ensured that all the produced bioconcrete could be quantified. With these modifications, we were able to greatly reduce the loss of the bioconcrete yield as seen in **Figure S1** where the amount recovered increased from ∼3.3 g/L to 6 g/L, the expected yield. At the same time, we improved the reproducibility of our results by reducing the variation among biological replicates. This is important because reproducibility is crucial when the quantity produced is being used as a measure to distinguish various bacterial species in order to identify top bioconcrete formers.

We used an environmental sample (seawater) as inoculum to identify high bioconcrete-producing consortia, then isolated top performers from the consortia that precipitated the most bioconcrete, all without including urea. Our approach varies from the recent literature. Many recent studies focus on isolating bacteria first (generally with urea in the medium), testing them for ureolytic activity, and, typically, then choosing the most ureolytic isolates to test for bioconcrete-forming abilities (2,23,34–42). Some deviations from this include Kakad and Shede’s study in 2023 performing an enrichment culture in media containing calcium before isolating the bacteria from their samples, and Ekprasert et al. in 2020 isolating non-ureolytic bacteria by streaking to isolation colonies that made bioconcrete crystals (43,44). Enrichment cultures that allow early elimination of non- or low-producing bioconcrete formers, such as what we present here, may allow more expedient isolation of better-performing bacteria from various environments. Additionally, prior researchers studying bioconcrete production in marine environments designed an artificial seawater recipe with expensive ingredients (45,46); our media used here, which are modifications of the commercially available Marine Broth medium, are inexpensive and straightforward to produce.

Due to the higher yield of bioconcrete by the sea isolates *B. sp.* SBS7 and *M. sp.* SBS5 in comparison to the other non-ureolytic bacteria studied, these isolates may be more promising candidates for bioconcrete application in the sea environment than the commonly studied *S. pasteurii* and *Ahb. pseudofirmus*. Due to its ability to sporulate and its higher viable cell count at the end of the 16 day experiment, *B. sp.* SBS7 is especially promising, as sporulation will allow it to withstand the stresses of incorporation into a concrete structure as well as lie dormant until the structure cracks; also, with more viable cells, there likely can be more healing. As the field of marine bioconcrete develops, a safety framework is needed to establish the suitability of different seawater bacterial isolates for industrial and field applications. Of note, the non-ureolytic control species used here, *B. toyonensis*, has a history of detailed safety evaluation because of its use as an animal feed additive (Toyocerin®; 47–49).

*S. pasteurii* produces bioconcrete very quickly owing to its high ureolytic activity. Conversely, the non-ureolytic organisms presented here produced bioconcrete on a relatively much slower scale - 16 days to the 1 day of *S. pasteurii*. The speed of production can impact the application for which each of these bacteria may be suited for. Slower production may allow for deeper crack healing and typically has the benefit of not requiring urea or producing ammonia. Faster healing may be better for sealing the surface quickly to prevent the ingress of harmful ions, but at the expense of requiring urea and potentially poor healing of deeper cracks. For application of bioconcrete to self-healing of a submerged, artificial coral reef substrate, we posit that the slower mechanism would prove for better healing and longevity, without requiring urea that may negatively impact the reef colonizers. This requires further investigation.

Our sea isolates form bioconcrete on the timeline of weeks rather than days, and we did not enrich for ureolytic or denitrification activity while culturing them. Carbonic anhydrases are nearly ubiquitous in cellular organisms, but vary in quantity and activity across species (50). We hypothesize that our seawater isolates use their carbonic anhydrase activities to produce bioconcrete in the culture conditions used here. Browsing their genome annotations in NCBI shows that all three seawater isolates are predicted to encode at least two carbonic anhydrases, with SBS7 predicted to encode four. Future work will be required to determine the exact mechanism(s) underlying the urea-free production of bioconcrete by these isolates. It follows that with the bioconcrete enrichment protocol detailed here, we may actually be enriching for isolates with higher carbonic anhydrase activity (and/or calcium binding ability) over others in the consortium. This higher activity may be due to the presence of more carbonic anhydrase genes/classes of carbonic anhydrase, higher transcriptional activity, faster enzymes/substrate turnover, or a combination of these. We propose that further study of these factors may enable the discovery of useful strains for carbon-negative, urea-free bioconcrete production.

## Acknowledgements

This material is based upon work supported by the U.S. National Science Foundation under Grant No. 2318123. Any opinions, findings, and conclusions or recommendations expressed in this material are those of the author(s) and do not necessarily reflect the views of the National Science Foundation. This work was additionally supported by the Cecil H. and Ida Green Chair in Systems Biology Science to K.P. We gratefully acknowledge Dr. Matthew Ramsey for providing the seawater used in our experiments. We gratefully acknowledge the Olympus Discovery Center and the Imaging Core facility at UT-Dallas for providing equipment and support.

---

**Figure S1:**
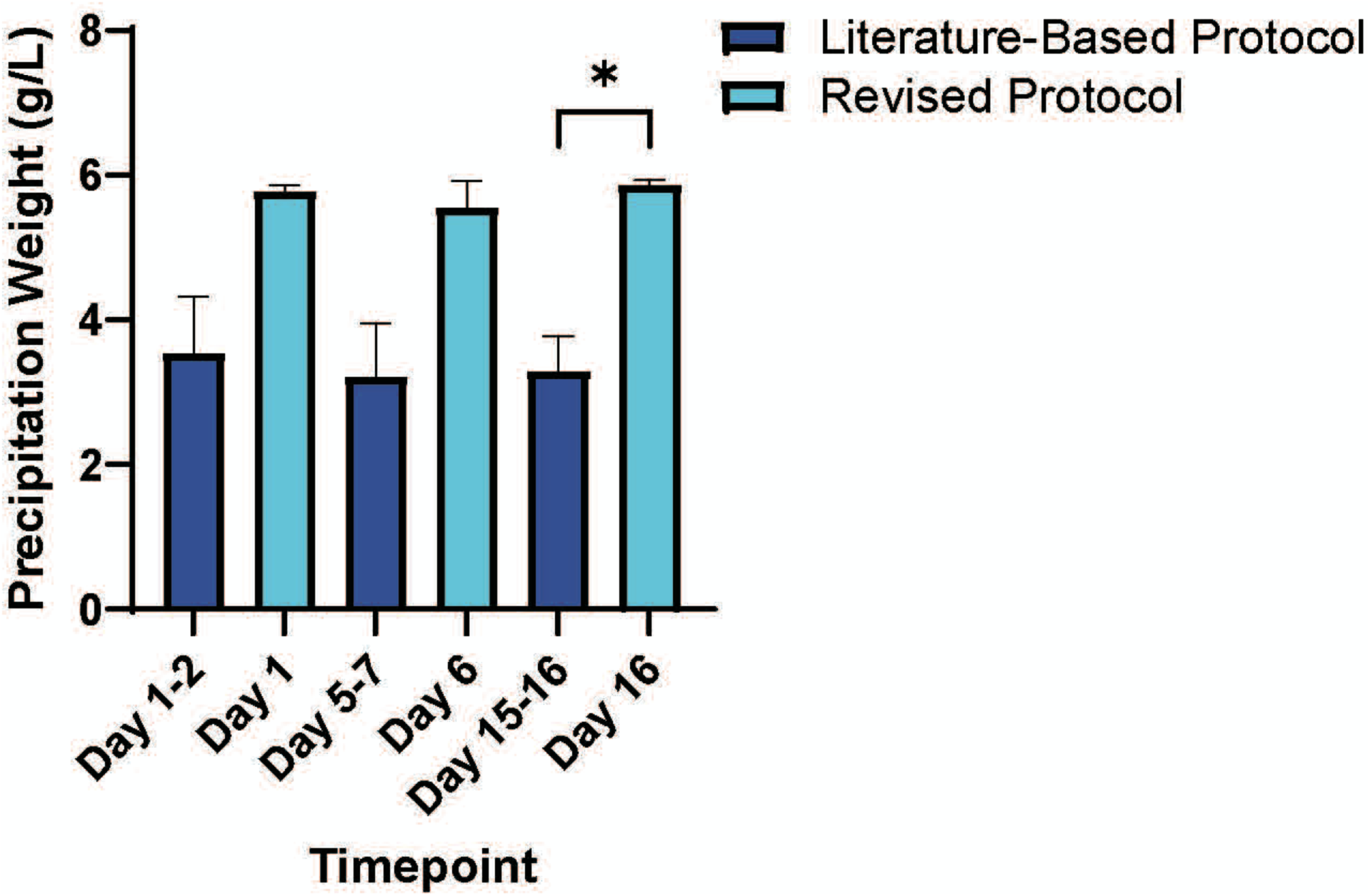
Amount of bioconcrete precipitated over time by *S. pasteurii* in LBCaU media using the literature-based and revised protocols. Friedman tests on the data for each individual protocol are not significant indicating no significant difference among timepoints, consistent with *S. pasteurii’s* fast production. A one-tailed Mann-Whitney test comparing the amounts produced by the different protocols by the day 16 timepoint shows a significant difference (p=0.05) in the amount of bioconcrete recovered indicating the new protocol improves the amount collected.

**Table S1.** Manufacturer and catalog numbers of chemicals used in media from this study.

| **Chemical** | **Manufacturer** | **Catalog Number** |
| --- | --- | --- |
| Tryptone | Fisher BioReagents | BP1421-500 |
| Yeast Extract | Bacto | 212750 |
| Sodium Chloride | Fisher BioReagents | BP358-212 |
| Instant Ocean Sea Salt | Instant Ocean Spectrum Brands | SS3-50 |
| Instant Ocean Reef Crystals Reef Salt | Instant Ocean Spectrum Brands | RC3-50 |
| Urea | Fisher BioReagents | BP169-500 |
| Calcium acetate hydrate, 99% | Alfa Aesar | A14947 |

**Table S2.**
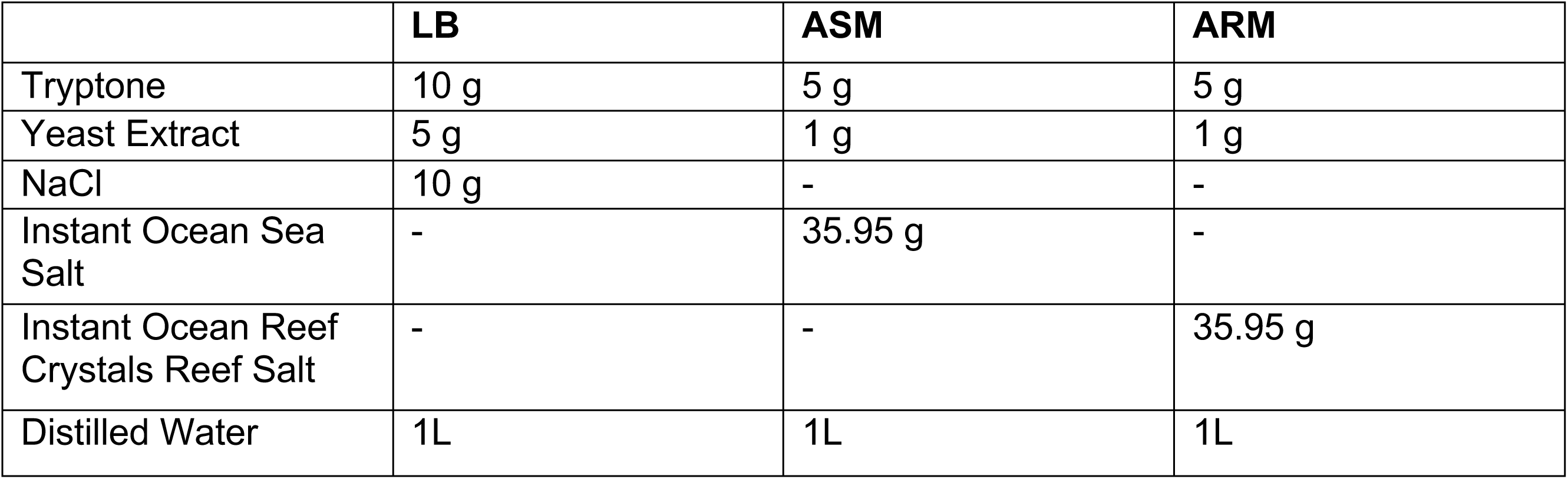
Comparison of ingredients in the base media used for bioconcrete quantification.

**Table S3.**
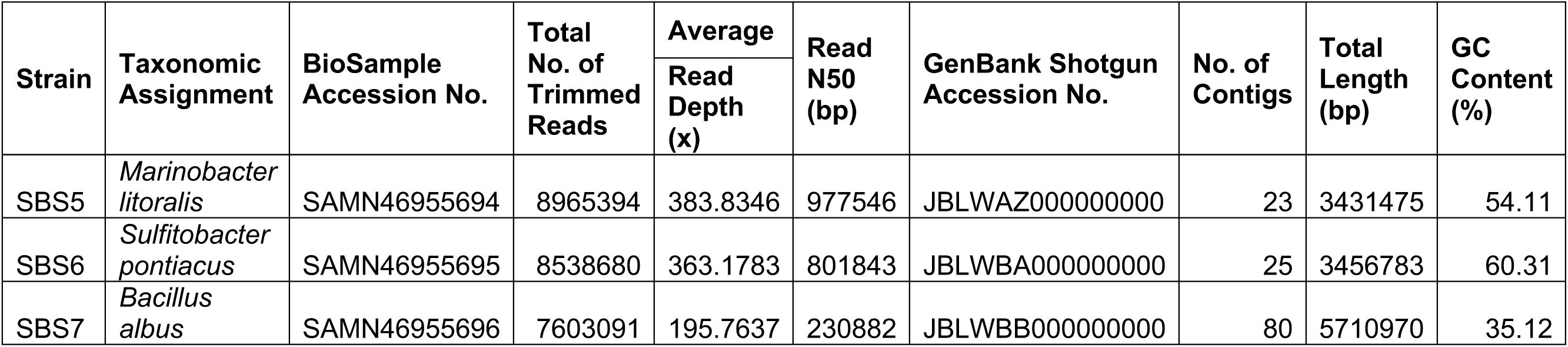
Features of the sea isolates’ genome assemblies.

**Supplemental File S1.** General bioconcrete quantification protocol for 1 (or 3) timepoint(s), modified from Reeksting et al, 2020, *Applied and Environmental Microbiology* (https://doi.org/10.1128/AEM.02739-19).

1. Streak bacteria on appropriate agar medium.
2. Use a single colony to inoculate overnight broth cultures.
3. Inoculate 15 mL medium in 50 mL centrifuge tube to achieve a starting OD_600_=0.001. If analyzing multiple timepoints, inoculate 50 mL media to achieve a starting OD_600_=0.001 in a glass flask and incubate 2 hours before aliquoting 15 mL into three 50 mL centrifuge tubes.

a. Completely close lids of centrifuge tubes, mark straight down the lidtiside, then open half a turn (mark on lid now rotated 180° from mark on side) and tape in place to ensure consistent oxygenation of cultures.
4. Incubate until the timepoint(s).
5. Remove one tube from the incubator at the timepoint for quantification.
6. Vortex to mix thoroughly.
7. Use 30 μL of culture to test pH via pH strip
8. Do 1:10 serial dilutions of culture, plate them on appropriate agar medium, and incubate to calculate CFU/mL.
9. Centrifuge the remaining culture at 780 x g for 4 minutes.
10. Aspirate the supernatant.
11. Wash with 40 mL autoclaved water.
12. Aspirate the supernatant.
13. Repeat wash and aspiration steps 2 more times (for 3 total washes).
14. Pre-weigh 1.5 mL microcentrifuge tube.
15. Resuspend bioconcrete pellet using 1 mL autoclaved water and transfer using wide-bore pipeAetip to pre-weighted 1.5 mL microcentrifuge tube

a. Cucng the tip of a normal pipeAe tip achieves a wide-bore tip.
16. Cover tube with Kim wipe and incubate at 55°C for at least 2 days
17. Remove tube from incubator, uncover and close tube, weigh.
18. Subtract pre-weighted tube amount to calculate the weight of the bioconcrete.

